# G-Trap Assay I: Small GTPases as sensitive immune response biomarkers for active bacteremia

**DOI:** 10.1101/2022.06.09.495550

**Authors:** Ross Clark, Virginie Bondu, Peter Simons, Laura Shevy, Stephen Young, Eliseo F. Castillo, Nancy L. Kanagy, Angela Wandinger-Ness, Thomas Howdieshell, Tione Buranda

## Abstract

The annual toll of sepsis is a 33% mortality rate for hospitalized patients with a cost of greater than 60 billion dollars in the U.S. There is a correlation between sepsis mortality rates and time to treatment with broad-spectrum antibiotics. Consequently, antibiotics are prescribed to nearly every patient suspected of bacteremia. However, once determined that broad-spectrum antibiotics are not required, it is unclear how to optimize the de-escalation of the antibiotics. There is an urgent need for methods to distinguish bacteremia from sterile inflammation and to assess antibiotic efficacy. Rho (Rac1 and RhoA) and Ras (Rap1) family GTPases are dynamic nodes of signaling convergence used by immune-activated leukocytes migrating to sites of infection. This study targeted the onset of GTPase activation as a biomarker of infection-induced immune activation in trauma patients. GTP binding assays were performed using a novel GTPase effector trap flow cytometry assay (G-Trap). Here, we demonstrate increased GTP binding to small GTPases (Rac1, Rap1, and RhoA) of resting cells serially exposed to plasma samples from bacteremic trauma patients. Responses to the serial samples showed that GTPase activation was influenced by the concentration of circulating pro- and anti-inflammatory mediators, in tandem with synergy or antagonism from the cytokines and the antibiotic treatment.

## INTRODUCTION

Sepsis is a life-threatening organ dysfunction syndrome caused by a dysregulated host response to infection.[1] Sepsis is a significant source of long-term morbidity and the cause of 30-50% of in-hospital deaths in the United States.[2] In addition, the financial burden comprises 40% of all pre-COVID-19 ICU expenditure, with total costs exceeding $60 billion.^[3–5]^ Bacterial, viral, fungal, or parasitic infections may develop into sepsis and can accompany non-infectious etiologies, making diagnosis challenging when an infection source is ambiguous.[6] Blood cultures remain the standard and most widely used method to assess for bacteremia. However, several days are required for positive culture samples if they manifest. Each hour of delayed clinical intervention in a patient presenting with a presumptive focal or bloodstream infection is associated with an increased risk of death.^1^ Therefore, patients presenting for emergency medical care are prescribed empiric broad-spectrum antibiotics before culture results become available. However, inappropriate prescription rates and overuse of antibiotics undermine antibiotic stewardship efforts.[7]

Alternatives to blood cultures such as the SeptiFast™ multiplex PCR kit (Roche) and IRIDICA™ (Abbott Diagnostics),[8] introduced a few years ago as potential rapid bacteremia detection platforms, are yet to gain commercial viability. Newer diagnostic technologies on the market include the FDA-cleared T2Candida, and T2 bacteria DNA panels for detecting pathogens in under 4 hours (www.t2biosystems.com) or ePlex, a PCR-based system for detection and identification of Grampositive (GP), Gram-negative (GN), and fungal pathogens as well as antimicrobial resistance genes within 1.5 hours using positive blood culture following gram staining. While promising, the drawback of these systems is reliance on blood culture growth. The detection of certain species cannot differentiate a positive result as a pathogen or a skin contaminant. More significantly, because of tissue localization of infection or prophylactic initiation of patient treatment with antibiotics, nearly 70% of blood culture results are negative.[9–12] More recently, measuring the host response to infection has emerged as a promising strategy. Using host immune mRNA panels integrated with machine learning algorithms to accurately identify the presence and type of infection (virus, bacteria, fungus) anywhere in the body has been described in pre-FDA-approved protocols. [6, 9]

Cytokines are essential to innate and adaptive immune responses whereby activated T helper cells (T_h_ cells) and macrophages become the primary cytokine producers. Cytokine receptor engagement elicits intracellular signaling cascades leading to altered gene expression in target cells, which leads to the trafficking of leukocytes to sites of infection.[13–15] The host response to infection is initiated when a pathogen is recognized by the innate immune cells such as neutrophils, macrophages, and dendritic cells, including some epithelial cells endowed with toll-like receptors that recognize evolution-conserved pathogen-associated molecular patterns on pathogens.

Depending on the host-pathogen risk factor assessment, proinflammatory cytokines such as tumor necrosis factor (TNF-a), interleukin (IL) 1, and IL-6 are secreted to stimulate the production and recruitment of neutrophils and macrophages. In addition, the small GTPases of the Rho family (Rac1, RhoA)[16–18], and Ras family (Rap1) GTPases [19–21] are critical regulators of cellular actomyosin dynamics, which drive leukocyte migration.[17, 22–26] The activation of Rac1, Rap 1, and RhoA depends on the exchange of GDP for GTP that is facilitated by guanine nucleotide exchange factors (GEFs) in response to chemotactic factors and cytokine binding to cell surface receptors.[27–30] Receptor occupancy induces GEF-mediated GTP binding to GTPases[31–34] (**Fig. 1**).

**Figure 1.**
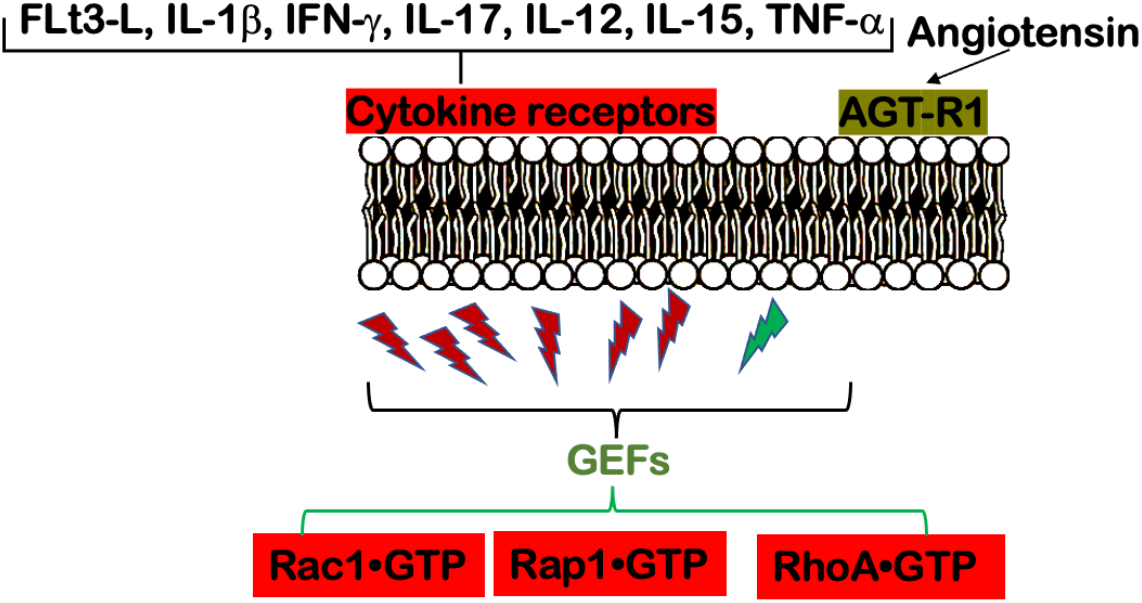
Schematic of cell membrane receptor-mediated GTP-binding to Rho GTPases. Rho GTPases cycle between active GTP-bound and inactive GDP-bound forms. The activity of Rho GTPases is governed by the availability of GAPs (GTPase-activating proteins), GEFs (guanine-nucleotide-exchange factors), and GDIs (guanosine-nucleotide-dissociation inhibitors). The activation of small GTPases such as Rac1, Rap 1, and RhoA depends on the exchange of GDP for GTP that is facilitated by guanine nucleotide exchange factors (GEFs). The conformation of GTP-bound GTPase results in specific binding to effector proteins (such as the PAK1, Ral, and Rhotekin for Rac1, Rap1, and RhoA, respectively), which do not recognize GDP-bound protein. Of relevance to this study, infection leading to sepsis elicits the release of various inflammatory mediators and angiotensin binding to cognate receptors that initiate signaling pathways that increase the intracellular concentration of GEFs. As shown in the schematic, GTP binding to small GTPases involves the convergence of multiple signals related to immune response. We devised an assay called G-Trap that can quantitatively measure activated GTP-bound Rho family GTPases in response to host immune-inflammatory mediators. The assay can be used to test the immune functionality of leukocytes directly.

This study used serial blood plasma samples collected from patients with confirmed bloodstream infections to stimulate GTP binding to Rac1, Rap1, and RhoA in quiescent cells. Our goal was to test whether the composition of pro- and anti-inflammatory mediators in the plasma of infected patients significantly impacted the magnitude of GTPase activation in target cells and thus establish small GTPases as immune integrative[22] biomarkers of infection. In addition, we evaluated the correlations between GTPase activation and host cytokines expressed in serial plasma samples from trauma patients, including those diagnosed with bloodstream infections.

## RESULTS

### G-Trap assay selectively measures GTP-bound GTPases

The activation of small GTPases such as Rac1, Rap 1, and RhoA depends on the exchange of GDP for GTP that is facilitated by guanine nucleotide exchange factors (GEFs) in response to chemotactic factors binding to cell surface receptors.[27–30] The conformation of GTP-bound GTPase results in specific activation of downstream effector proteins (such as the PAK1, Ral, and Rhotekin for Rac1, Rap1, and RhoA, respectively) (**Fig. 2A**). [35–37] We devised an assay called G-Trap[37] capable of quantitatively measuring activated GTP-bound Rho family GTPases due to host immune-inflammatory mediators and demonstrating value for early recognition of patient immune response to infection. [37] For initial proof-of-principle, we used lipopolysaccharide (LPS), a bacterial endotoxin, plasma from a patient with a culture-confirmed infection, and Fms-like tyrosine kinase-3 ligand (Flt-3L), a hematopoietic growth factor, [38, 39] to stimulate GTP binding to Rho GTPases in phagocytes and endothelial cells via a common pathway[40] (**Fig. 2B-D**).

**Figure 2.**
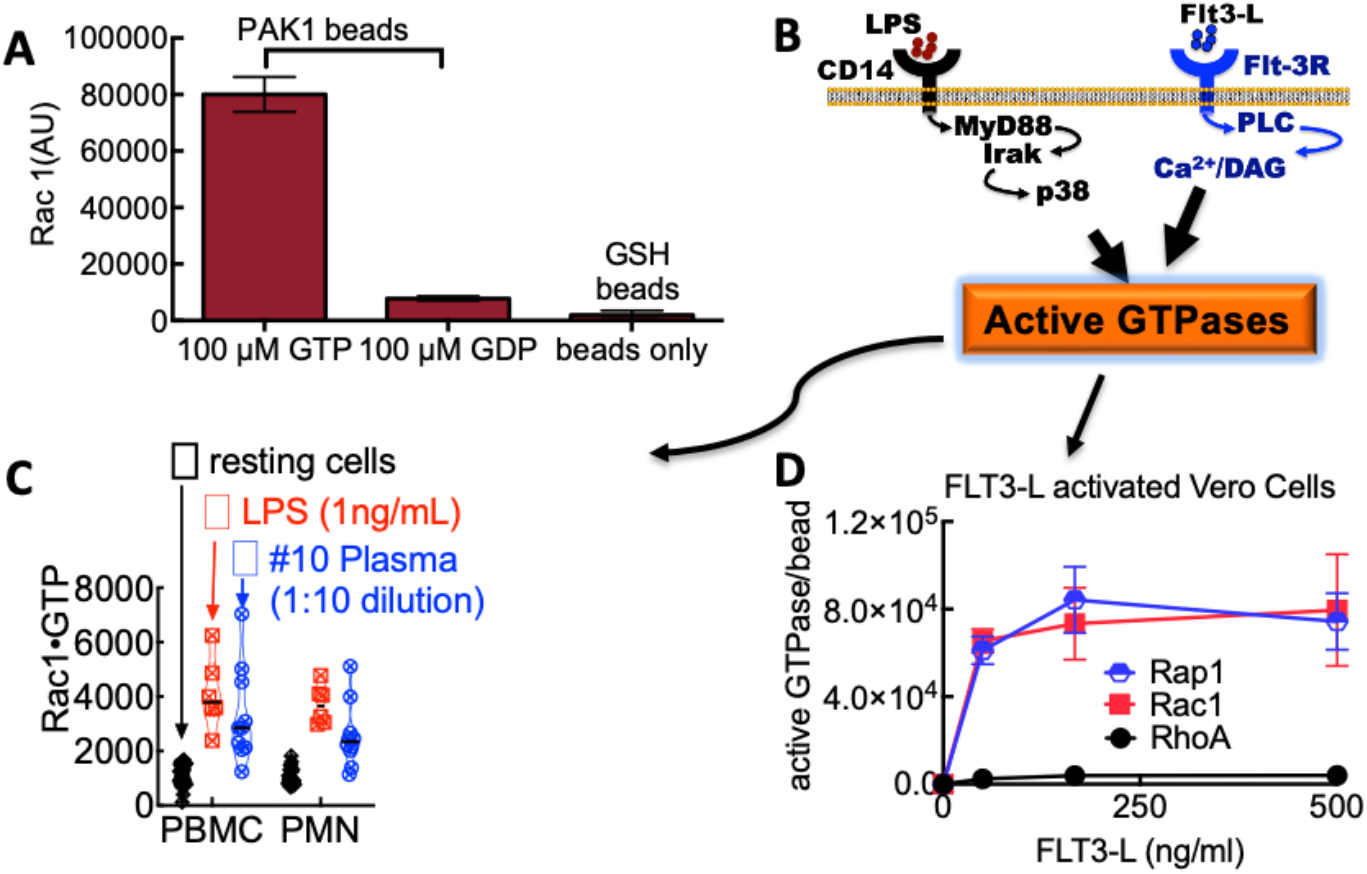
Principles of G-Trap assay functionality. The assay uses multiplex beads functionalized with p21 activated kinase protein binding domain (PAK-1 PBD), a Rac1 effector, Rhotekin-RBD, a Rho effector protein, RalGDS-RBD, a RAP1 effector protein. The conformation of GTP-bound GTPase results in specific binding to effector proteins, which do not recognize GDP-bound protein. **A**. Relative binding of Rac1-GTP and Rac1-GDP to GST-PAK1-RBD functionalized beads. **B**. Endothelial and immune cells express receptors for recognizing pathogen-associated molecular patterns (PAMPs) such as LPS from bacterial cell walls. Also, the host expresses cognate receptors of inflammatory mediators. Occupancy of immune receptors initiates signaling required to activate Rho family GTPases. Model of bacterial and Flt-3L cytokine-induced signaling pathways leading to activation of small GTPases (e.g., Rac1, Rap1, and RhoA) inside infection-fighting leukocytes. **C**. Stimulated GTP binding to Rac1 in peripheral blood monocytes (PBMCs) and polymorphonuclear leukocytes (PMN) from uninfected controls is upregulated by exposure to lipopolysaccharide (LPS) or plasma from a patient (Patient #10 in this study) with confirmed bacteremia (Haemophilus influenzae). **D**. 30 min exposure to Flt-3L induced GTP binding to Rap1 and Rac1 but not RhoA in Vero cells. 50,000 cells in 48 well plates were incubated with (0, 50, 167 and 500 ng/ml) Flt-3L for 30 min at 37°C. Cell lysates were analyzed for GTPase activity.

### GTPase activating cytokines circulate in the plasma of patients with confirmed bacterial infection, demonstrating a correlation between cytokine expression and GTPase activity

#### a. Cytokine expression

The immune system is designed to recognize a wide variety of pathogens. The immune response is believed to be partitioned into three types (1, 2, and 3) established to antagonize different classes of pathogens. Type 1 responses are initiated by the production of IL-12, which primarily elicits the release of IFN-γ, a cytokine designed to eliminate intracellular threats such as viruses and bacteria. Type 2 reactions deploy IL-4, IL-5, and IL-13 against large extracellular parasites, and type 3 responses are primarily initiated by IL-1β and IL-23 to produce effector IL-17 and IL-22 as effector cytokines devised for extracellular bacteria and fungi.[41] In addition, the immune system includes an anti-inflammatory component (T regulatory cells, T_regs_) designed to regulate the pro-inflammatory branch of the immune response, thus preventing severe pathogenesis. [42] In **Fig. 3**, we compare the relative magnitudes of type 1, 2, and 3 responses of two trauma patients, Patient #3, who suffered terminal sepsis, and Patient #18, who survived multiple nosocomial infections. The data points span the patients’ lengths of stay at UNMH. In general, peak cytokine expressions for Patient #3 were greater than for Patient #18. Still, the overall differences between patients and controls were not substantial, except for Patient #3’s expression of IFN-γ, IL-17, and IL-1RA, which was significantly higher than Patient #18. Type 2 cytokine expression was comparable between patients and controls.

**Figure 3.**
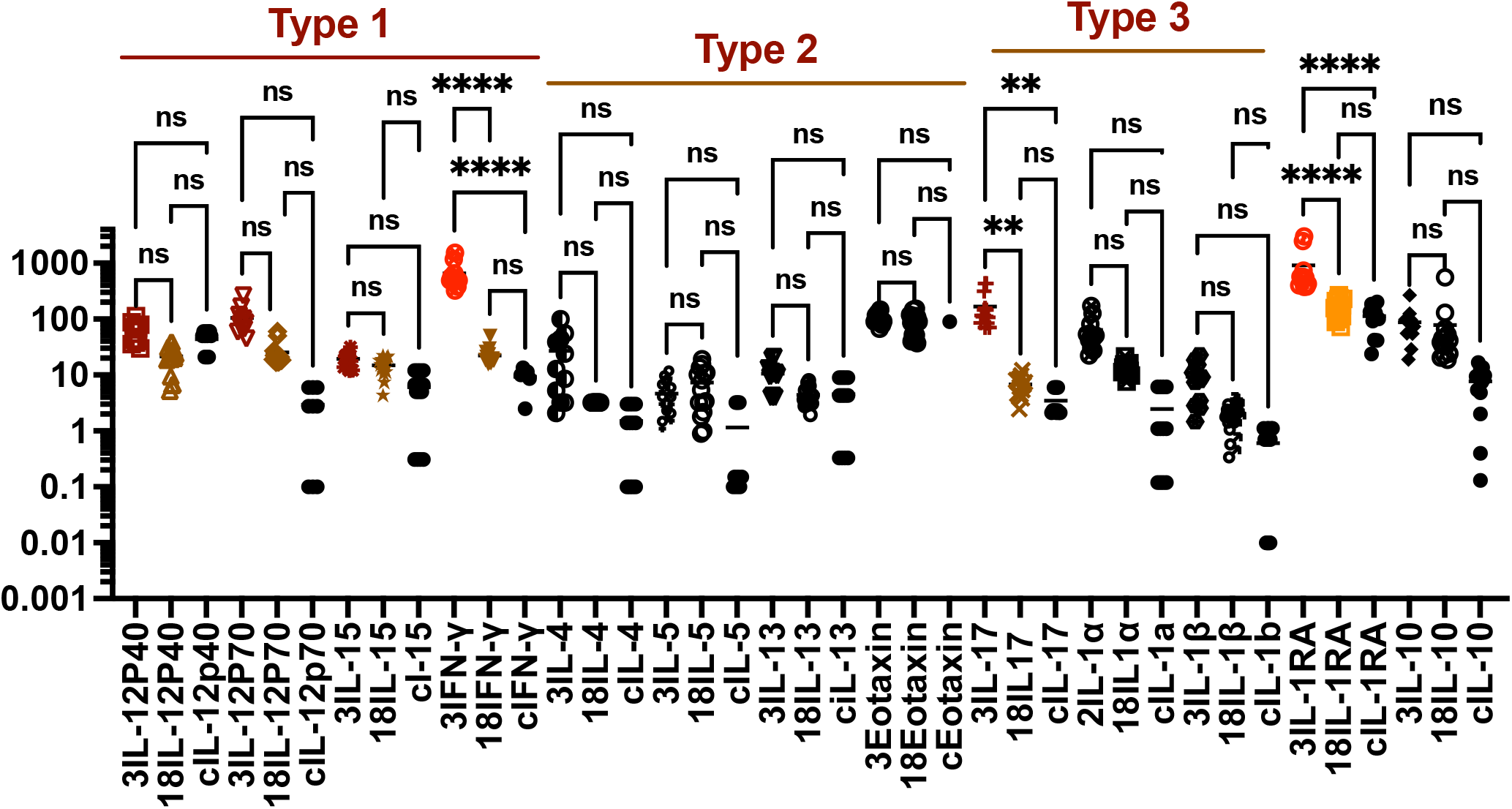
Relative magnitudes of type1, 2, and 3 immune responses of patients #3 and #18. Traditionally type 1 responses are against intracellular pathogens (e.g., viruses and bacteria), type 2 against extracellular pathogens ((e.g., helminths), and type 3 against extracellular organisms (e.g., bacteria and fungi).

#### b. Cytokines induce GTPase activation

We next examined temporal relationships between the expression of Flt3-L, type 1 and 3 cytokines, and GTPase activity in serial samples of Patients #3 and #18. We superimposed normalized cytokine levels and active GTPases to visualize correlation trends (**Fig. 4**). *<u>Patient #3’s</u>* immune response oscillated between peaks and troughs (days 2, 6, and 9 in **Fig. 4A-B**). Due to extensive trauma injuries, the patient required endotracheal intubation and mechanical ventilation on admission. The initial hospital course indicated systemic inflammatory response syndrome (SIRS) and infection. On day 2 the patient developed signs of infection, including leukocytosis (supplementary **Fig. S1**) and >100-fold increase in C-reactive protein (CRP), a known marker of sepsis. Interestingly, peak expressions of IL-1RA and IL-17 switched relative positions at the top on days 2 and 6, respectively. In contrast, pro-inflammatory IL-12p70 and IFN-γ expression in serial samples persisted in the top 3 cytokines correlated to GTPase expression. Cultures of blood and sputum were obtained, and sputum culture was positive for *Streptococcus pneumoniae on day 6*. The overall immune response declined as the patient developed progressive multiple organ dysfunction, including worsening ventilatory requirement and hepatic failure. The patient expired on post-injury day 10.

**Figure 4.**
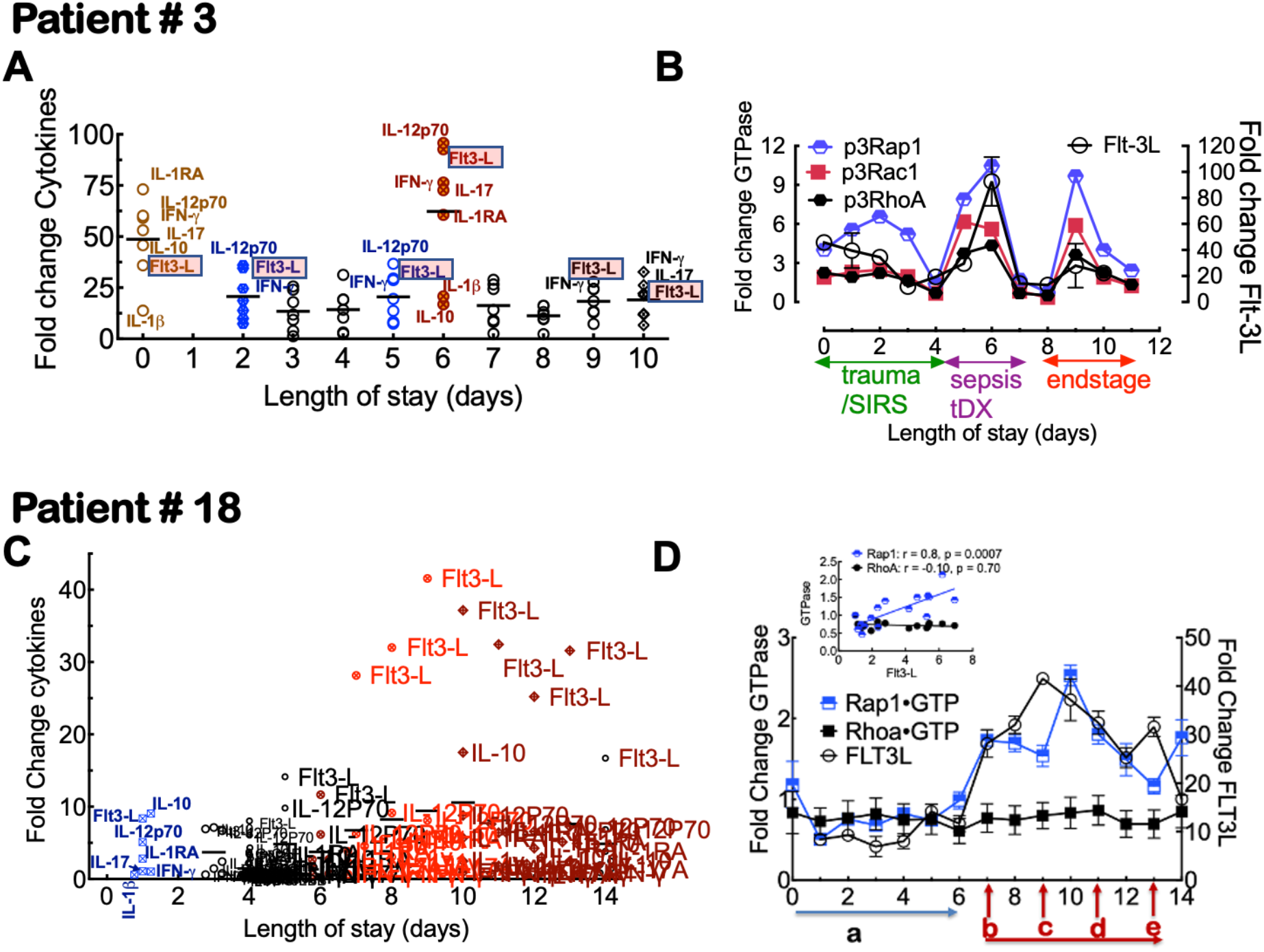
<u>*Patient #3*</u>, an 88-year-old patient, was a pedestrian in a motor vehicle collision. **A**. Plot of Flt3-L, type 1 and type 3 cytokines. **B**. Overlay of Rap1·GTP Rac1·GTP, RhoA·GTP, and Flt-3L showed overlapping peak expressions at different time points. Firstly, day 2 coincided with the clinical need for crystalloid infusion. Secondly, Flt3-L peaked on day 6, when the patient’s blood culture was determined to be positive for the bacterial pathogen. Thirdly, Flt3-L peaked two days after a blood transfusion following refractory hypotension. *Patient #18*_was a 50-year-old patient admitted after a motor vehicle collision, suffering severe injuries, including bilateral pulmonary contusions and multiple rib and vertebral fractures. **C**. Analysis of the longitudinal expression of 41 cytokines (HCYTMAG-60K-PX41 Cytokine kit, Human; Millipore, MA) showed that FLT3-L was significantly elevated above other cytokines on consecutive days that preceded clinically confirmed infection. **D**. GTP binding to Rap1 and RhoA (in Vero E6 cells), superimposed with Flt3-L expression. Notably, the rise in FLT3-L and Rap-1 activation occurred 48 h before the positive blood culture (tdx) was drawn. However, within 36 h after antibiotic treatment was inaugurated, FLT3-L and Rap1 activation began to decline following the patient’s response to treatment. **Insert** Spearman rank correlation indicates a positive correlation between Rap1-GTP and Flt3-L expression. **a**. patient on mechanical ventilation. **b**. Percutaneous tracheostomy **c**. febrile, ventilator-associated pneumonia (VAP), empiric antibiotics (Vancomycin, Zosyn) later positive for methicillin-sensitive staph aureus (MSSA). **d**. Atrial fibrillation (beta-blockade and amiodarone treatment). **e**. day 13 and 15, additional blood cultures drawn, later positive for staph epidermis, patient prescribed two-week course of Vancomycin.

*<u>Patient #18</u>*, admitted after a motor vehicle collision, suffered severe injuries. The patient required mechanical ventilation on admission. ICU length of stay (LOS) was 16 days. The first 6 days were characterized by immune homeostasis[43] (**Fig. 4C**), indicating the relative expression levels of type 1, 3, and immunoregulatory cytokines. Post day 6, the patient’s immune response was dominated by Flt3-L, which drove GTPase activity (**Fig. 4C**) as shown above (**Fig. 2D**). The patient developed central line-associated bloodstream infection (CLABSI) and became febrile on day 7/8. Blood cultures drawn on day 9 were positive for methicillin-sensitive *Staphylococcus aureus* (MSSA). The patient was treated with empiric parenteral vancomycin and piperacillin-tazobactam. Additional blood cultures drawn on post-injury day 11 were positive for *Staphylococcus epidermidis* and *Staphylococcus aureus* and accompanying tracheobronchitis. The patient received a complete two-week course of vancomycin ending on post-injury day 25.

#### c. Correlated Cytokine release regulates GTPase activation

We next evaluated pairwise Pearson’s correlations of Flt3-L, Rap1, Rac1, RhoA, type 1, 2, and 3 cytokines and IL-10, IL-1RA, IL-6, and IL-8. IL-6 and IL-8 are typically associated with neutrophil activation.[44] We used the breaks in cytokine and GTPase activity expression patterns to divide the data into three temporal segments (**Fig. 5**). For Patient #3, the cytokine correlations were generally robust throughout the patient’s length of stay, especially the middle segment, days 5-7, when the patient was culture-positive from an infection. In contrast, the correlation matrix of Patient #18 immune variables presented a relatively weak mixed inter-and intra-immune-type cohesion. Overall, the peak elevation of GTPase activity of Rap1 was increased 10-fold from baseline levels for Patient #3, whereas Patient #18’s immune response elicited a two-fold increase.

**Figure 5.**
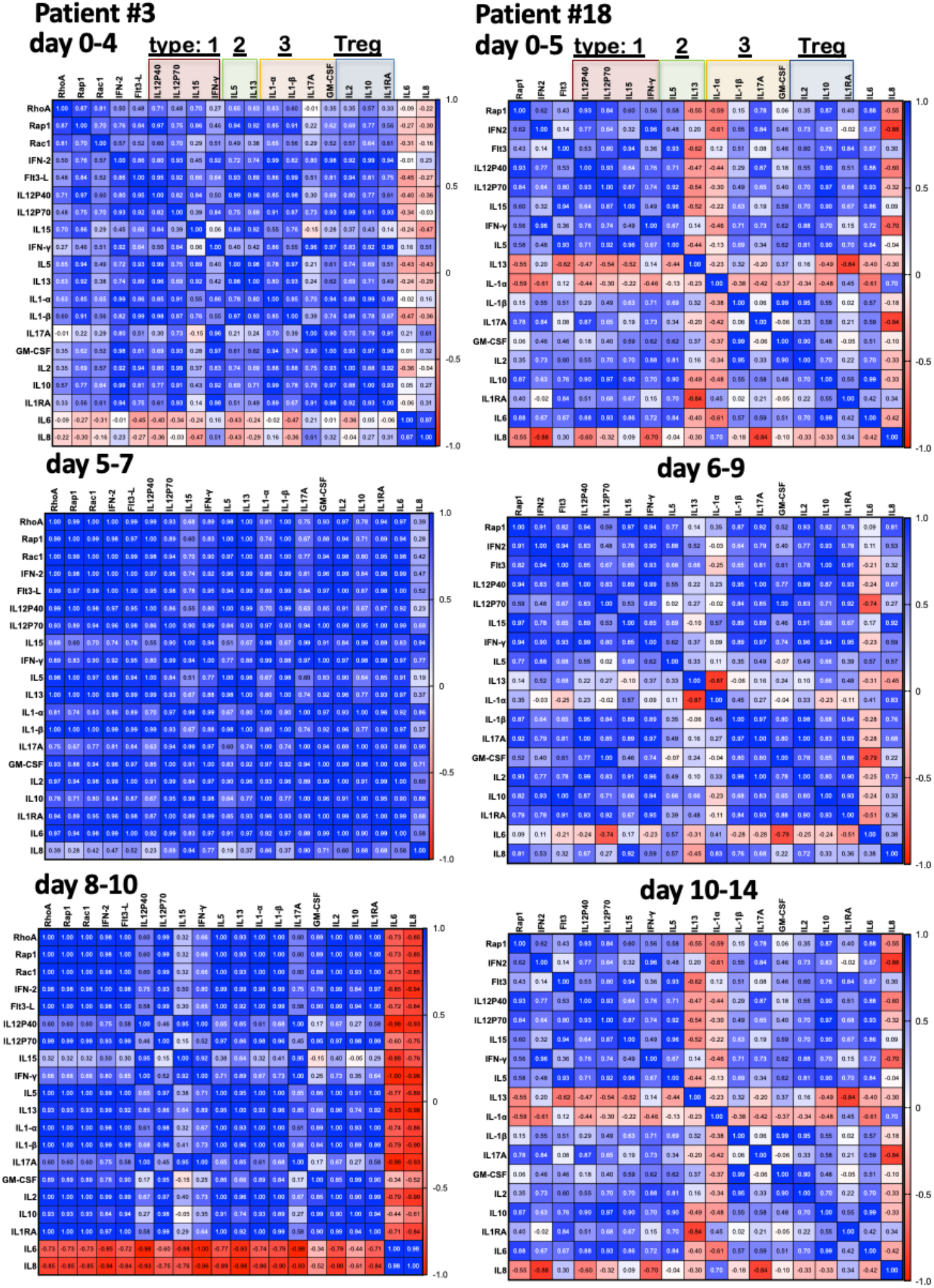
Pearson’s correlation matrices for GTPase activity and cytokine expression over time intervals are specified for each panel for patients #3 and #18. Based on the relative correlation strengths, patient #3’s immune mobilization is more robust than patient #18. Correlations are classified according to the following scale: 0–0.19 very weak, 0.2–0.39 weak, 0.4–0.59 moderate, 0.6–0.79 strong, 0.8–1.0 very strong. **D**. Pearson’s correlation matrix for the day 5 – 7day interval. **E**. Pearson’s correlation matrix for days 8-10.

#### d. Principal Component Analysis

The data are presented as a distance biplot of Principal component (PC) scores (patient samples) and *PC loadings* vectors (cytokines, active GTPases; Rac1, Rap1, and RhoA). The first PC (PC1) represents the most variance in the data. The second PC (PC2) is perpendicular to the first PC and represents the second most variance in the data axis. The *PC loadings* vectors indicate a proportionate concentration of the immunological variables and correlation with serial patient samples. In addition, proximity between *PC scores* and specific *PC loadings* vectors shows a strong correlation. Conversely, vectors appearing at ~180° are negatively correlated, and those at 90° are not correlated. In **Fig. 6A**, the first component shows the strong correlation of type1, type 3, FLT3-L, and GTPases which is reflected in the more remarkable fold changes in concentration of the cytokines, indicating the association of GTPase activity with pro-inflammatory immune response. On the other hand, the PC2 axis signified disposition to neutrophil activation with IL-6 and IL-8.[44] In addition, the correlated TNF-a/MCP-1 pair was reminiscent of inflammatory network response to sepsis and mortality.[45] For Patient #18, the distribution of variables in the first component shows a predisposition to a weakened immune response due to anti-inflammatory response indicated by correlated IL-2 and IL-17 vectors (at ~180°) opposite IL-6 as the main determinants of PC1 (**Fig. 6B**). [46, 47] Furthermore, on days 10 through13, patient #18’s presentation of an overt shift towards type 2 immune response, clustering IL-1RA, IL-12p40, IL-5, and IL-13 along PC2 could be related to immune compromise. [48] In addition, the contribution of type 1 cytokines to the PC2 component shows the dissemination of immuno-suppression as both type 1 stimulating cytokines IL-12p70 and IL-15 are positioned at a cosine angle of 135° to IFN-γ. Patient #18 was immunosuppressed to the extent that their type 1 effector cytokine response was comparable to non-infected trauma patients (Supplementary **Fig. S2**) and experienced opportunistic polymicrobial bacteremia.

**Fig. 6.**
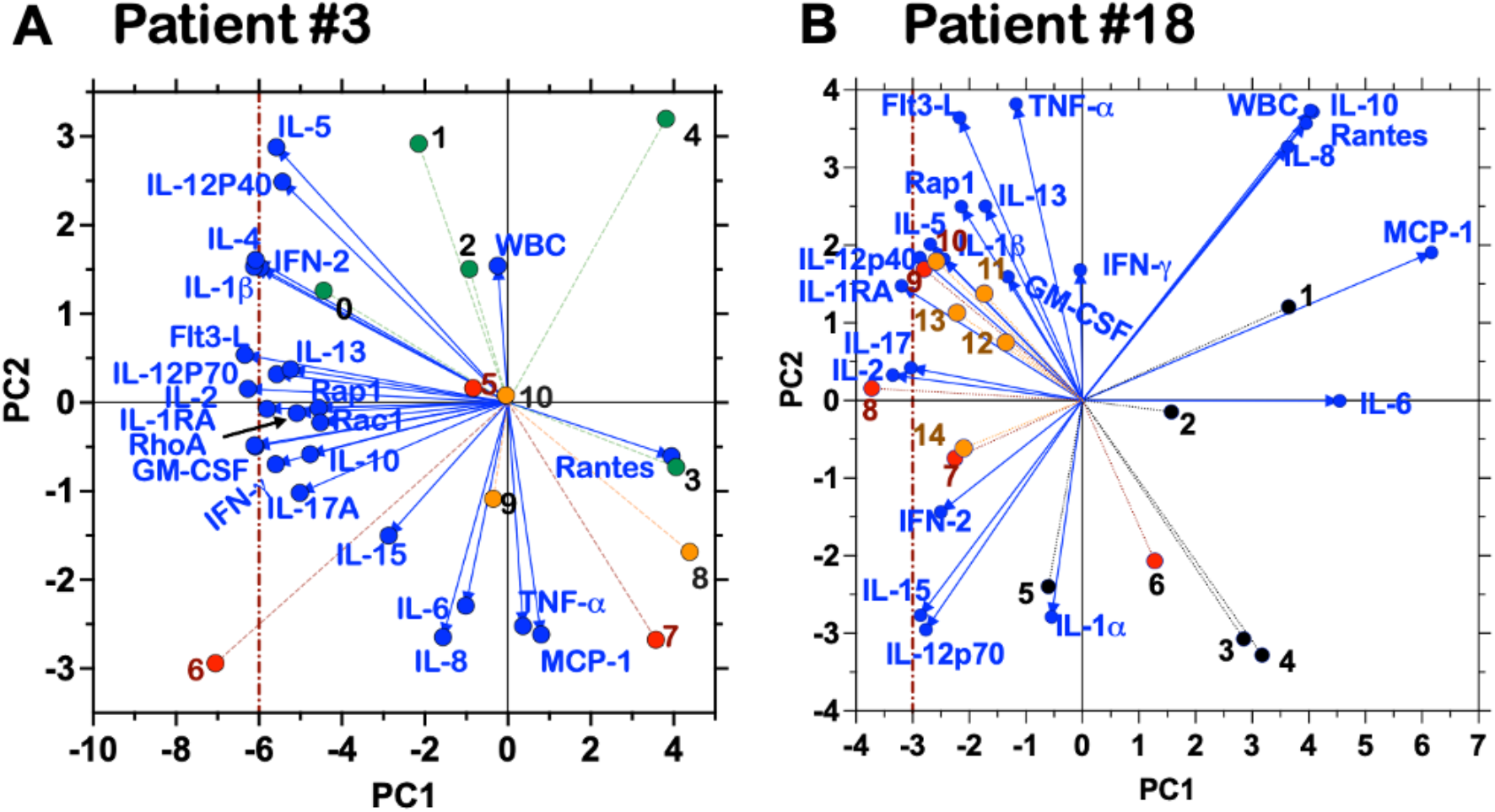
Principal component analysis (PCA)of the immune response of Patient #3 (**A**) and Patient #18 (**B**). The PCA biplot graphs comprise PC loading vectors represented by variables (Rap1, WBC, and innate and adaptive immune cytokines) and PC scores represent daily patient samples associated with the analytes. PC1 axis represents variables with the most variance and PC2 with the second most variance. The vertical dotted lines indicate the limit of PC1 for the two patients. The correlation coefficient between the variables is defined by the cosines of the angles between the loading vectors they represent, where an angle close to zero indicates a high correlation between variables, and a 90° angle shows no correlation. A 180° indicates a negative correlation. The vectors’ magnitude and the PC scores’ coordinates relative to the vectors indicate the weight of the mutual relationship between the two. Thus, in Panel A, the PC score 5 linked to the sample from day 5, carries minimal weight located near the origin of the graph, compared to PC score 6.

### GTPase activity is sensitive to antibiotic treatment failure in patients with a bacterial infection. Patient #10

Patient #10 was a 53-year-old individual involved in a motor vehicle collision and sustained injuries that included right hemopneumothorax, left pneumothorax, bilateral clavicle fractures, right scapula fracture, right 1-9 rib fractures, right flail chest, and left 1-7 rib pneumomediastinum, and bilateral L5 pars interarticularis fractures. ICU length of stay was 25 days. Patient #10 required mechanical ventilation and vasopressor support during the early part of the hospitalization. Bacterial infection was suspected on hospital day 2 due to the onset of fever and changes in vital signs (*e.g*., temperature, respiration rate, tachycardia) (**Fig. 7A**). Thus, parenteral vancomycin (*customarily prescribed to treat serious, life-threatening infections by Gram-positive bacteria unresponsive to other antibiotics*) and piperacillin-tazobactam were initiated empirically. (*Tazobactam is a beta-lactamase inhibitor that makes piperacillin effective against organisms that express beta-lactamase and would generally cause degradation of beta-lactamase-sensitive antibiotics. However, tazobactam shows little antibacterial activity by itself and, for this reason, is not administered alone*). On hospital day 6, a sputum culture grew *Haemophilus influenzae*, and the antibiotic regimen was narrowed to ceftriaxone (*to which Haemophilus influenzae was susceptible in this case*). Antibiotics were stopped three days later on hospital day nine but resumed again on day 14 after clinical evidence of a relapse of the infection. Recrudescence of infection was corroborated by increased Rap1•GTP (5-fold) and Rac1•GTP (7-fold) levels over baseline levels on day ten. Thus, the G-trap assay results presaged the clinical need to resumption antibiotic treatment by three days (**Fig. 7B**). Ultimately, no other pathogen was isolated in blood or sputum cultures, though the patient improved with further antibiotic treatment. This clinical narrative demonstrates the utility of the G-Trap assay in assessing antibiotic effectiveness or infection persistence where existing clinical tools failed and necessitated empiric treatment.

**Figure 7.**
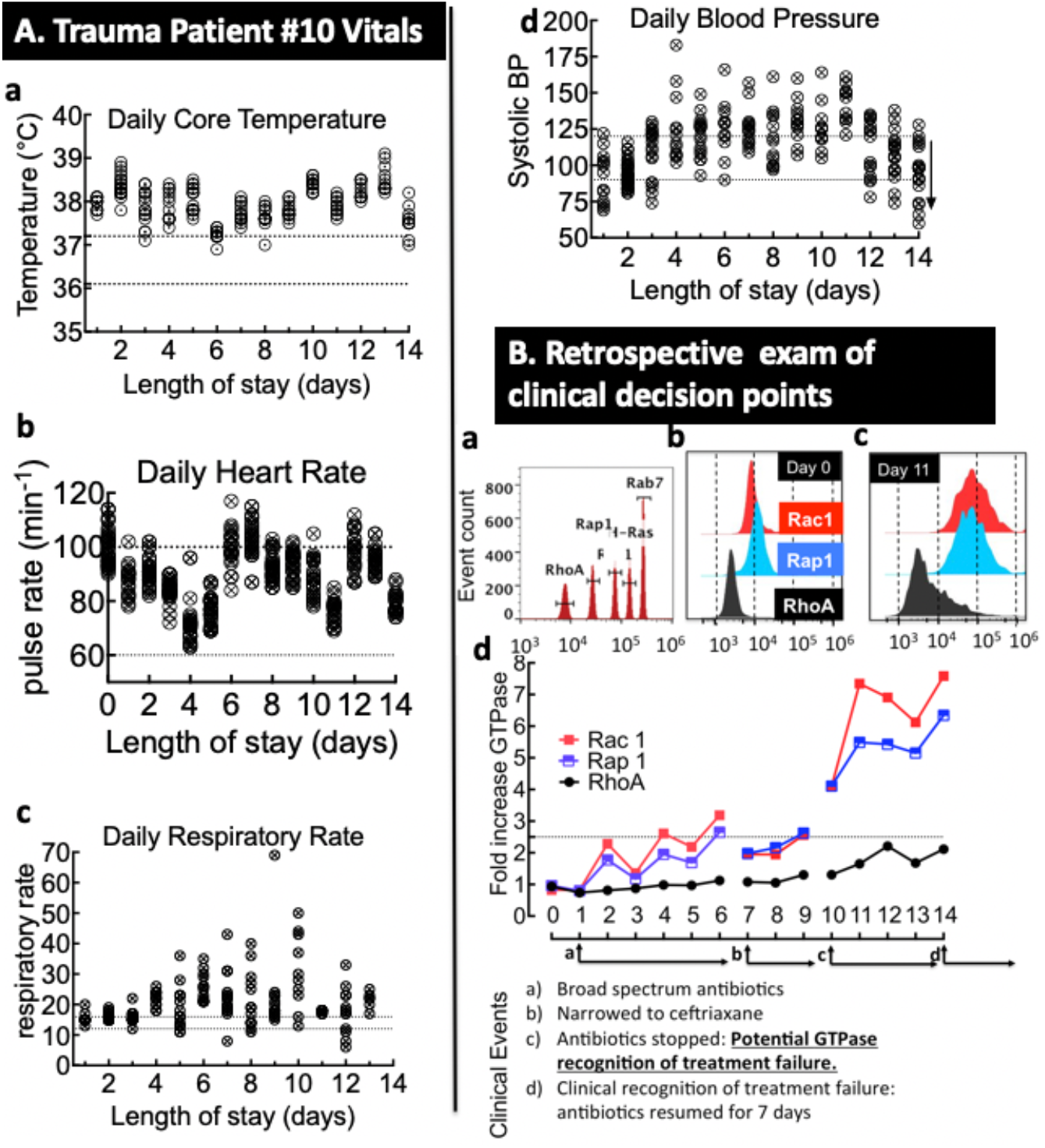
*Patient #10* was a 53-year-old individual involved in a motor vehicle collision. **A**. Panels show the patient’s vital signs. Dotted lines represent the normal range. **B**. G-Trap data readout, showing: **a**. Histograms of Cyto-Plex^™^ beads coded with discrete levels of 700 nm fluorescence beads for target GTPases. **b&c**. Flow cytometry histograms associated with GTPase activity in cell lysates exposed to blood plasma drawn from Patient #10 shown in **d** on days 0 (pre-infection) and 11 (post-infection). G-Trap assay measured Rap1-GTP and Rac1-GTP cells activated with serial samples and the clinical narrative.

### Proteomic Analysis of Blood Plasma from Infected Patients Reveals additional GTPase Activating Immune Factors

Proteomic analyses identified further potential contributors to the significant increases in GTP binding to Rap1 and Rac1 observed in patient #10. For these analyses, we examined the samples from days 8-11. Laser densitometry readings of the plasma samples were computer analyzed for differential expression of proteins as previously described.[49] 2D-Gel spot data showing significant differences between serial samples designated as controls were determined from adjusted p-values using FDR (and Bonferroni) analysis as outlined in the methods. We selected five spots from patient #10 for mass spectrometry analysis (**Fig 8A** and Table 1) based on Pearson’s correlations of gel spots and G-Trap data (**Fig. 8B**). Carboxy peptidase N (CPN) and T-cell receptor (TCR) β-chain were significantly correlated to Rap1 and Rac1 activation. Another marker, c-reactive protein (CRP), was strongly correlated to CPN.

**Figure 8.**
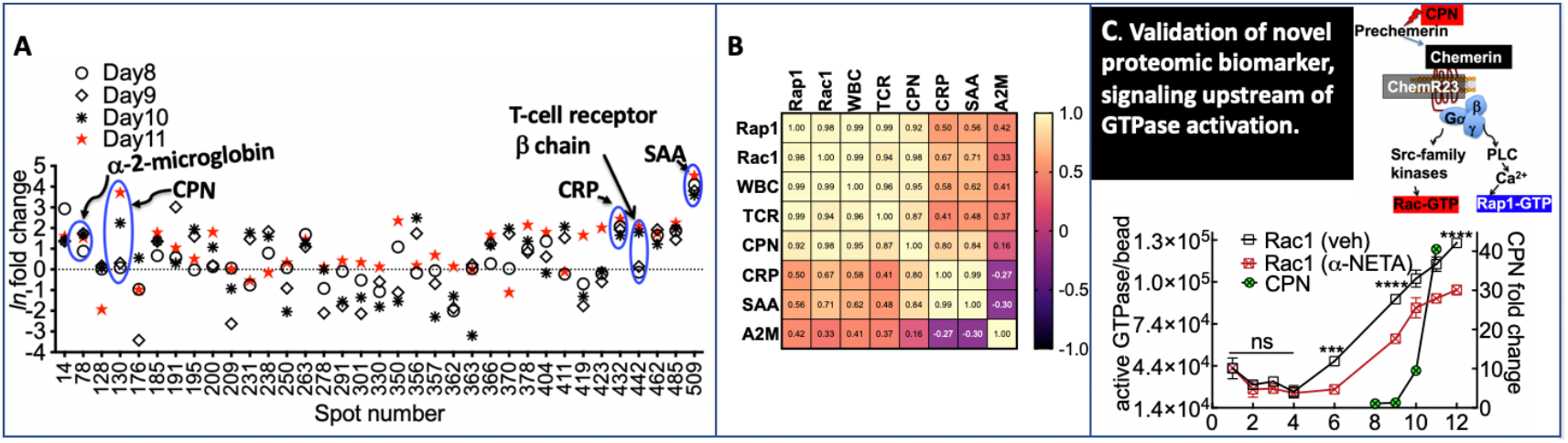
*Patient #10*. GTP loading to Rap1, Rac1, and RhoA, to Rho family GTPases is correlated to the up-regulation of bacterial infection biomarkers. **A**. Plot of the logarithm of geometric mean fold change relative to control for select 2D gel spots derived from the plasma analysis of Patient #10. Spot reference numbers are on the horizontal axis, and there is one value for each day of sample collection. The spots marked with blue circles are associated with statistically significant (False Discovery Rate) changes relative to control for the days indicated. The markers include, spot #78; a-2-microglobulin (A2M), spot #130; CPN, spot #432; C-reactive protein (CRP), spot #442; T-cell receptor ß chain (TCR) and spot #509; serum amyloid A (SAA). **B**. Pearson’s correlation for Patient #10 Rap1 and Rac1 versus TCR, CPN, A2M, CRP, and SAA. **C**. Proteomic analysis revealed *carboxypeptidase N (CPN*) as a potential *candidate activator of GTPase*. CPN is a metallocarboxypeptidase that regulates complement anaphylatoxins (C3a, C4a, and C5a). CPN expression is superimposed over changes in GTP binding to Rac1 and Rap1 for days 8-11. CPN exerts its proteolytic activity on prochemerin, a zymogen for chemerin, a potent chemoattractant *to its receptor ChemR23 (CMKLR1), a GaiPCR* for adaptive and innate immune responses recognized by ChemR23 receptor and induces GTP binding to Rap1 and Rac1 downstream of ChemR23. We used a-NETA, an antagonist for ChemR23 (K_d_ ~ 6 μM), and confirmed the presence of CPN-activated chemerin in Patient #10’s plasma. The data show partial sensitivity to a-NETA; suggesting that multiple factors such as cytokines contribute to GTPase activation after day 6.

The expression profile of CPN on days 8, 9, 10, and 11 were correlated to both Rac1 and Rap1 activity (Pearson correlation: ρ = 0.98, p<0.01) (Fig. 6C). CPN is a metallocarboxypeptidase that regulates complement anaphylatoxins (C3a, C4a, and C5a).[50–52] For a mechanistic rationale for selecting CPN (**Fig. 8C**), we reasoned that CPN exerts its proteolytic activity on prochemerin, a zymogen for chemerin, that is a potent chemoattractant (K_d_ ~ 0.1 nM)[53] and binds to receptor ChemR23 (CMKLR1). Very recent studies, published after our research was completed, have identified chemerin as a potential biomarker for the early onset of sepsis in critically ill patients.[54] Therefore, we used a-NETA, an antagonist for ChemR23 (*K_d_* ~ 6 μM), to test for the presumptive presence of CPN-activated chemerin in patient #10’s plasma. As shown in **Fig. 8C**, measurements of Rac1-GTP in cells challenged with the patient’s plasma from patient #10 on days 2-4 suggest that ChemR23 was not activated. However, on day 6, the data show partial sensitivity to a-NETA, suggesting an escalation of the inflammatory response once antibiotic use was inappropriately terminated early. The partial inhibition of GTP-binding to Rac1 induced by a-NETA is consistent with the idea CPN was one of several inflammatory mediators present in the plasma of patient #10, including cytokines, as indicated for patients #3 and #18.

### GTPase Activity is Sensitive to the Physiological Response to Hypotension. Patient #14

Patient # 14 was a 53-year-old individual who suffered minimal injuries in a single-vehicle collision. The patient was suspected to be septic on arrival with community-acquired pneumonia, evidenced by hypotension (**Fig. 9A**). Signs of sepsis were present throughout the ICU stay. Empiric parenteral vancomycin and piperacillin-tazobactam were started 4 hours after admission to the ICU; the following day, azithromycin was added for atypical bacterial coverage in community-acquired pneumonia. On day 3, the piperacillin-tazobactam was changed to ceftriaxone targeting the culture-confirmed *Streptococcus pneumoniae*, and vancomycin and azithromycin were continued. On day 4, vancomycin was discontinued. On hospital day 5, the patient developed gastric perforation of unclear etiology, diagnosed as a missed injury at presentation (rather than stress-induced gastric ulceration) requiring laparotomy. Metronidazole was added to the treatment regimen of ceftriaxone and azithromycin. On day 6, azithromycin was discontinued. On day 7, ciprofloxacin was added. At this point, the patient was on ciprofloxacin, ceftriaxone, and metronidazole. Ciprofloxacin and metronidazole were continued for broader spectrum antimicrobial coverage in the setting of gastric perforation and potential intra-cavitary contamination. After seven days, ceftriaxone was discontinued as the patient had completed the planned treatment regimen for pneumonia.

**Figure 9A.**
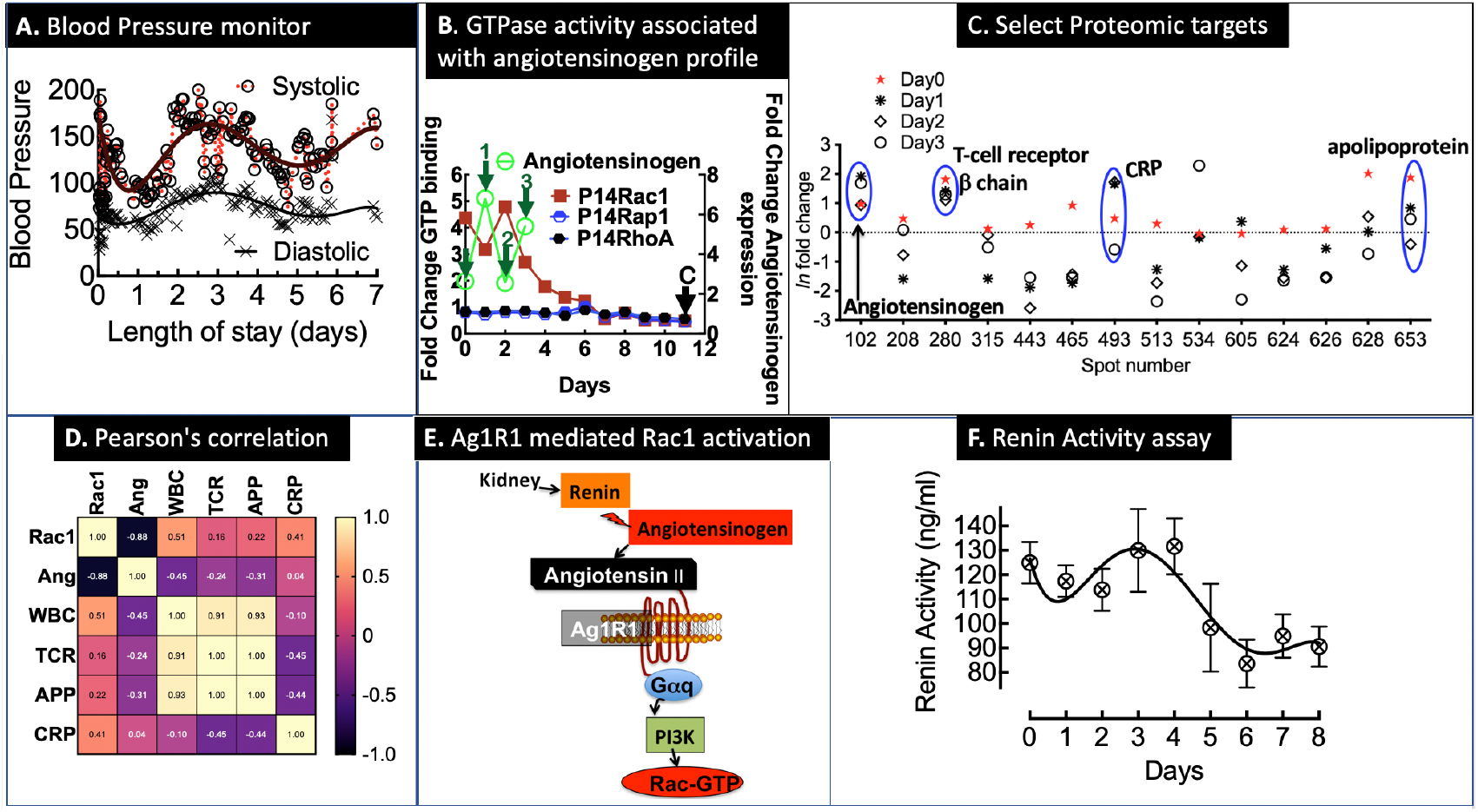
*<u>Patient # 14</u>*, a 53-year-old individual admitted with no injuries after a low-speed car collision. **B**. Overlay plot of fold-changes in GTP binding to Rac1, Rap1, and RhoA measured in cell lysates of Vero E6 cells after exposure to serial plasma samples (1:10 dilution) collected from trauma Patient #14. The letter C arrow denotes the control sample with baseline GTPase activity. Downward arrows on days 0-3 indicate samples analyzed for proteomic markers of GTPase activation. Angiotensinogen (Aogen) is a discovery target selected from four candidates based on unity with clinical data described in the text. We selected Aogen as the most probable and testable effector of GTPase activity relative to the other candidates, as shown in 5B and 5C. **C**. Plot of the logarithm of geometric mean fold change relative to control for select 2D gel spots derived from the plasma analysis of Patient #14. Spot reference numbers are on the horizontal axis, and there is one value for each day of sample collection. The spots marked with blue circles are associated with statistically significant (False Discovery Rate) changes relative to control for the days indicated. The markers include spot # 102; Aogen, spot # 280 (TCR), spot # 493 C-reactive protein (CRP), spot #653; Apolipoprotein (APP), **D**. Pearson correlation for Patient #14 Rac1 versus Aogen, TCR, CRP, and APP. **E**. Schematic of the mechanism of renin-induced Rac activation. Angiotensinogen is the preprohormone of angiotensin-II, a ligand for AgtR1, which induces vasoconstriction in a Rac1-dependent pathway. **F**. Plot of renin activity measured in serial plasma samples from Patient #14.

Serial plasma samples collected during the patient’s hospitalization were used to test for induction of GTPase activity in Vero E6 cells. Analysis of GTPase activation in cell lysates of Vero E6 cells exposed to patient plasma showed elevated GTP binding to Rac1 for the first four days that diminished to baseline levels in subsequent days. RhoA and Rap1 were inactive throughout the hospital stay (**Fig. 9B**). An analysis of the blood pressure readings and Rac1 activity was in sync with the temporal cycle of the sinusoidal blood pressure pattern of the patient (**Fig. 9A**). Targeted proteomic analysis of plasma samples from days 0, 1, 2, and 3 identified inflammatory markers whose expression levels corresponded to the GTPase activity pattern (**Fig. 9C**). Based on the oscillatory blood pressure recordings that appeared to track the changes in GTP-binding to Rac1 asynchronously, we selected spot # 102, identified by mass spectrometry as angiotensinogen, as a likely target (**Fig. 9D**). Angiotensinogen is the preprohormone of angiotensin-II, a ligand for AgtR1, which induces vasoconstriction in a Rac1-dependent manner (**Fig. 9E**).[55, 56] The renin-angiotensin-aldosterone system (RAAS) regulates blood pressure and fluid balance. In the setting of sepsis, RAAS is activated to mitigate the low arterial blood pressure associated with the disease. Hence the sinusoidal blood pressure patterns shown in Fig 6A. Activation of the RAAS is initiated by the release of the enzyme renin to convert angiotensinogen to the inactive prohormone angiotensin-I, which is then converted in the lungs by the angiotensin-converting enzyme to the active hormone, angiotensin II. To validate the selection of angiotensinogen, we used a Renin Assay kit (MAK157 from Sigma) to test for the activity of renin, an enzyme secreted by kidneys to promote the production of vasoconstrictor angiotensin II to mitigate low arterial blood pressure.[57–60] As shown in **Fig. 9F**, the renin activity results were correlated to Rac1 GTPase activity and the hypotensive episodes observed in the patient and with decreases in the substrate, angiotensinogen. Thus, the G-Trap assay results were validated by the proteomic discovery of angiotensinogen expression, patient blood pressure readings, and the renin activity assay.

## Discussion

The rapid assessment of indicators of bloodstream or tissue localized pathogens is essential for enabling appropriate treatment of present-on-admission or nosocomial infections in the ICU. Because of the significant health risk associated with sepsis, empirical broad-spectrum antimicrobial treatment is used before definitive microbial identification, associated with an increased risk of mortality, development of antimicrobial resistance, and high medical cost. To narrow the temporal gap of uncertainty in evaluating the presence of infection, we developed the G-Trap assay to assess the requisite functional activity of the host immune response to an infection. The premise of the G-Trap assay is based on the idea that an active immune response to infection involves the release of many inflammatory mediators that initiate signaling cascades upstream of small GTPases that play a significant role as effectors of leukocyte trafficking, cell adhesion, and extravasation to tissue sites of infection. In this proof-of-concept study, we examined changes in GTP binding to Rac1, Rap1 and RhoA extracted from lysed cells after being exposed to plasma samples from trauma patients who experienced infection after admission to UNMH Level One Trauma center. Plasma samples from healthy controls and uninfected trauma patients produced similar results, consistent with a quiescent GTPase status in exposed cells. In contrast, infections caused 2 and 10-fold increases above the baseline of GTP-bound GTPases in samples from two patients presenting a compromised immune response and robust pro-inflammatory outcome, respectively.

The significant findings from the analysis of samples from a patient (#3), presenting a robust proinflammatory response, were that the convergence of type1 and type3 response cytokines induced robust downstream GTPase signaling. IL-17, which is known to stimulate GTPase activation; [61, 62] was strongly correlated to IL-12, GM-CSF, IFN-γ, IL-1β, and TNF-a upstream of a robust GTP loading to Rac1, Rap1, and RhoA. Excessive levels of IL-17, such as in Patient # 3 (peak expression on day 6 was 433.0 pg/ml; compared to controls 8.7 ± 11.7 pg/ml[63]) are comparable to or exceeding levels detected in plasma and tissues during sepsis resulting in multiple organ damage and death.[64] On the contrary, patient #18’s cytokine response showed the results consistent with immune impairment, first evidenced by the instances of nosocomial infections in the clinical setting, which were corroborated by a lack of IL-12 mediated T_h_1 immunity. [65] Upregulation of Flt3-L expression increases regulatory T cell (Treg) numbers[66, 67]. Thus, the dominance of Flt3-L expression relative to other cytokines such as IL-17 (peak level of 12.2 pg/ml on day 8) was consistent with patient 18’s weakened immune response, thereby setting the stage for an increased risk of opportunistic infections.[68, 69] Taken together, cytokine activity upstream of GTPase activating GEFs (Fig. 1) produced differential results based on the nature of the patient’s immune response.

We also analyzed plasma samples for additional GTPase activating biomarkers other than cytokines. We demonstrated a positive correlation between infectious disease severity and patient production of chemerin (patient #10) or renin (patient #14). Chemerin is an adipokine that acts as a pleiotropic chemotactic factor for macrophages, natural killer cells, and immature dendritic cells, capable of enhancing inflammation through recruitment and retention of macrophages at the sites of inflammation (reviewed in ref[54]) and alternately inhibiting neutrophil recruitment and secretion of pro-inflammatory mediators.[70] A recent prospective observational study found that chemerin was significantly higher at the onset of severe sepsis than in healthy controls. [54] Interestingly, the same study found a significant positive correlation between chemerin and other known biomarkers of early sepsis (CRP, procalcitonin). In patient #10, we matched the expression pattern in inflammatory mediators to the proteomic discovery of CRP and chemerin-mediated immune response, which supports the utility of the G-Trap assay as a tracker of immune functionality during an infection. As evident in patient #14, the progression of sepsis to septic shock leads to low blood pressure.[57–60] Accordingly, the results from the G-Trap assay recapitulated the RAAS functionality, in contrast to the cytokine-enriched milieu of the early sepsis stage of #3, #10, and #18.

The main strength of our study is establishing the concept that Rho-GTPases capture the convergence of multiple signaling cascades downstream of cognate receptors of inflammatory mediators. However, using plasma samples to activate culture cells and not allowing the patient’s direct assessment of immune cell functionality is a limitation of our study. We have initiated further work targeting the immune-performance status of infected patients’ leukocytes to assess the functionality of leukocytes in infected patients and the net effect of empiric antibiotic treatment.[71]

## MATERIALS AND METHODS

### Effector Proteins

Glutathione *S*-transferase (GST) tagged-effector chimeras used for the studies are as follows: p21 activated kinase protein binding domain (PAK-1 PBD), a Rac1 effector, was obtained from Millipore Sigma, Rhotekin-RBD, a Rho effector protein, was purchased from Cytoskeleton (Denver, CO), RalGDS-RBD, a Rap1 effector protein expressed and purified from a plasmid, kindly provided by Dr. Burridge (UNC-Chapel Hill) [72], and GST-RILP was prepared as previously described. [36] a-NETA was purchased from ENZO, Santa Cruz, CA.

### Antibodies

Monoclonal rabbit anti-Rap1 was obtained from Thermofisher, monoclonal mouse antibodies anti-Rho (A, B, C) clone 55, and the secondary antibody goat anti-mouse IgG (H+L) conjugated to Alexa Fluor 488 were obtained from Millipore Sigma. Monoclonal Rac1 antibodies (Cat. # ARC03) were purchased from Cytoskeleton.

### Buffers

We used the following buffers: a) 2X RIPA buffer; 100 mM Tris titrated with HCl to pH 7.4, 300 mM NaCl, 2 mM EDTA, 2 mM NaF, 2 mM Na_3_VO_4_, 2% NP-40, 0.5% sodium deoxycholate, and just before adding to the culture medium, 2 mM PMSF and 2X protease inhibitors. b) HHB buffer; 7.98 g/L HEPES (Na salt), 6.43 g/L NaCl, 0.75 g/L KCl, 0.095 g/L MgCl_2_ and 1.802 g/L glucose. HPSMT buffer, an intracellular mimic: 30 mM HEPES, pH 7.4, 140 mM KCl, 12 mM NaCl, 0.8 mM MgCl_2_, 0.01% Tween-20.

### Cell Culture

Vero E6 cells from the American Type Culture Collection (ATCC) were plated at 2×10^4^ cells per well of a 48-well plate, then allowed to grow for ~48 hours to medium/low confluence (50,000 cells/well) in Dulbecco’s Modified Eagle Medium (DMEM) supplemented with 10% fetal bovine serum (FBS), 100 units/mL penicillin and streptomycin, and 2 mM glutamine in a 37 °C incubator with 5% CO_2_. Telomerase Immortalized Microvasculature Endothelium (TIME) cells from ATCC were plated at 20,000 cells per well in 48 well plates and used the next day in low serum media. The cell density doubles overnight.

### Serial plasma samples from trauma patients

The study population (UNM HSC IRB #13-312) for plasma samples comprised trauma patients admitted to the University of New Mexico’s (UNM) Level 1 Trauma Center with bacteremia upon or acquired an infection during their hospitalization for post-injury care. The study was authorized to collect up to 14 plasma samples on consecutive days. The study recruited 20 patients, the patient population comprised 8 women and 10 men with severe trauma. The mean age was 49 (range, 24-90) years. However, most patients were disenrolled for inability to draw blood. In addition, only six patients were diagnosed with an infection. For this study, four patients (#s3, 10, 14, and 18) presented clinical evidence of bacteremia with vital signs and laboratory abnormalities consistent with sepsis and positive blood culture samples. We also analyzed plasma samples from two patients admitted with penetrating trauma (patients #17 and #19) who did not develop an infection.

Some samples were analyzed for cytokine expression and used to stimulate Vero E6 or endothelial cells in 48 well plates, lysed, and analyzed for changes in GTP binding to Rac1, Rap1, and RhoA. We compared the expression of GTP-bound GTPases to baseline levels of non-infected control subjects. We analyzed data seeking correlations between GTPase activation and cytokine expression. To rationalize cytokine-complementary mechanisms of GTPase activation, we carried out a proteomic analysis of serial plasma samples to identify inflammatory mediators stimulating GTP loading to Rap1, Rac1, and RhoA in exposed cells. To determine patient responsiveness to antibiotic treatment, we analyzed the changes in levels of GTP binding as a measure of the efficacy or failure of empiric antibiotic therapy.

### Plasma cytokines and chemokines

Cytokines were measured in duplicate plasma samples using an HCYTMAG-60K-PX41 Premixed 41 Plex - Immunology Multiplex Assay (Millipore). The results were subsequently analyzed using GraphPad Prism Version 9.2.0 software (San Diego, CA, USA). We used pooled healthy control samples (N = 20) for baseline representation.

### Rap1, Rac1, and RhoA GTP binding assays

GTP binding assays were performed using a GTPase effector trap flow cytometry assay (G-Trap). [36, 37] This protocol has been fully described in a methods chapter. [37] A 48-well plate is seeded with 20,000 Vero E6 or TIME cells in 100 μL of culture medium per well, resulting in about 50,000 cells the next day. First, the culture medium is removed and replaced with 100 μL of serum-free medium overnight. Next, 10μL of trauma patient plasma was added to wells and incubated at 37°C in an incubator. Next, the plate was chilled in an ice/water bath to conserve GTPase activity. Finally, each well was treated with 100μL of cold 2X RIPA buffer to lyse the cells. After centrifugation in a microfuge tube, the 200 μL of cleared lysate was used for triplicate tests.

### Targeted Proteomic Analysis of Patient Plasma Samples

*2D Gel Loading and Sample Preparation*. As previously described,*[49]* two-dimensional electrophoreses were performed by Kendrick Labs, Inc. (Madison, WI, USA); duplicate 2D gels were scanned with a laser densitometer (Model PDSI, Molecular Dynamics Inc., Sunnyvale, CA, USA).

#### Data Analysis

Replicate 2D gels were analyzed to generate 510 and 610 spot-intensity data for Patients #10 and #14, respectively (summary reports can be found in Supplementary data). In addition, laser densitometer readings of duplicate patient plasma samples (#10 and #14) were analyzed for differential expression of proteins. *[49]* Spot data showing significant differences between selected serial samples and controls were determined from adjusted p-values using false discovery rate (FDR) analysis. In addition, we used G-Trap results to select target plasma samples collected on four consecutive days where GTPase activity was determined to undergo dynamic changes relative to control samples. *[49]* Nine spots identified this way were selected for Matrix-Assisted Laser Desorption/Ionization (MALDI) mass spectrometry at Clarkson University Core facility.

### Renin Activity Assay

We used a Fluorometric Renin Activity assay kit (Ab138875) from Abcam to measure renin activity in the plasma samples of patient #14. The assay was set up according to the manufacturer-supplied protocol using a 384 well plate. However, we modified the buffer solution to include 2% Tween-20 detergent. Samples were analyzed using a Biotek H2 Synergy Plate Reader.

### Statistical Analysis

Data shown are presented as mean values ± SD. We used GraphPad Prism 9.3.1 for statistical analysis, where statistical significance was determined using paired two-tailed Student *t*-test or group-wise, one-way ANOVA analyses followed by multiple-testing correction using the Tukey method; *\*p* < 0.05, \*\**p* < 0.01, \*\*\**p* < 0.005, \*\*\*\**p* < 0.001. Two tailed Pearson’s pairwise correlations and principal component analyses were also performed using GraphPad Prism, Version 9.3.1 (**https://www.graphpad.com/scientific-software/prism/**

## Acknowledgments

This research was funded by NIH National Center for Research Resources, the National Center for Advancing Translational Sciences: UL1TR001449, and the UNM Department of Pathology.

---

**Figure S1.**
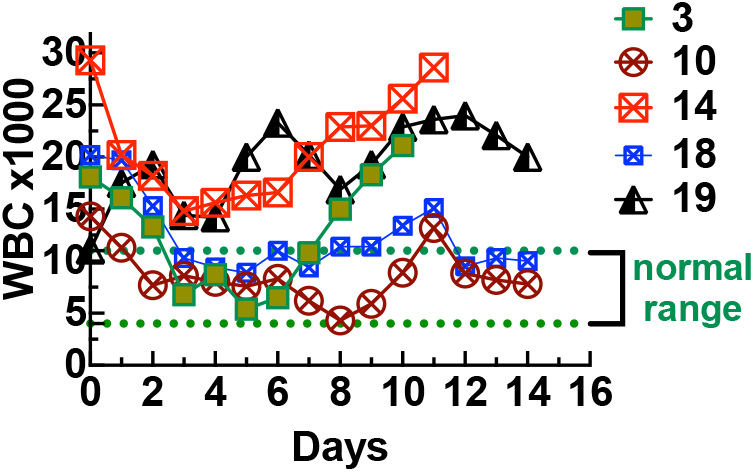
A plot of serial white blood cell (WBC) counts for case study patients.

## GTPase activity is proportional to the regulation of proinflammatory immune response

In inpatient #3, the immune response oscillated between highs and lows, representing one of the hallmarks of severe sepsis.[73] Patient #18 was immunosuppressed to the extent that they experienced a second infection with *Staphylococcus epidermidis* while on antibiotics for MSSA. To assess the immune functionality of the two patients, we next examined the temporal relationships between the pro-inflammatory cytokines IL-12(p40/p70) and IFN-γ.[74] IL-12 bridges the innate and adaptive immune responses and skews T-cell reactivity toward the production of IFN-γ production. p40 is a presumptive natural competitive receptor-binding antagonist of the biologically active p70. Consequently, the p40 subunit is secreted in ≥10 -fold excess over p70 in resting conditions.[74] Once the inflammatory immune response is initiated, the p40/p70 ratio is expected to decrease.

We compared the p40/p70 ratio and IFN-γ production of 20 control patient samples (pooled), two uninfected trauma patients with penetration injuries (#17 and #19), and patients #3 and #18. The results are summarized in **Figure S1**. For the controls, the p40/p70 ratio was 10:1, and IFN-γ was 18.0 pg/ml, which is comparable to the literature (15.4 ± 3.8).[75] The production of IFN-γ by patient #19 was similar to controls, whereas patient #17 was twice as high.

Patient #3’s temporal changes of p40/p70 fluctuated due to the imbalance in the production of IL-12 isoforms, where the peak expression of IFN-γ, Flt3-L (and GTP binding to Rap1 and Rac1 occurred on day 6 when the p40/P70 ratio was <1. However, temporal production of IFN-γ was 10-100-fold higher than controls and patient #18. This result indicates that patient#18’s impaired T helper 1 (Th1) response presaged the nosocomial infections on day 13, as noted above. The weaker immune response in patient #18 relative to patient #3 is reflected in the 1:5 ratio in their relative peak levels in GTPase activity.

**Figure S2.**
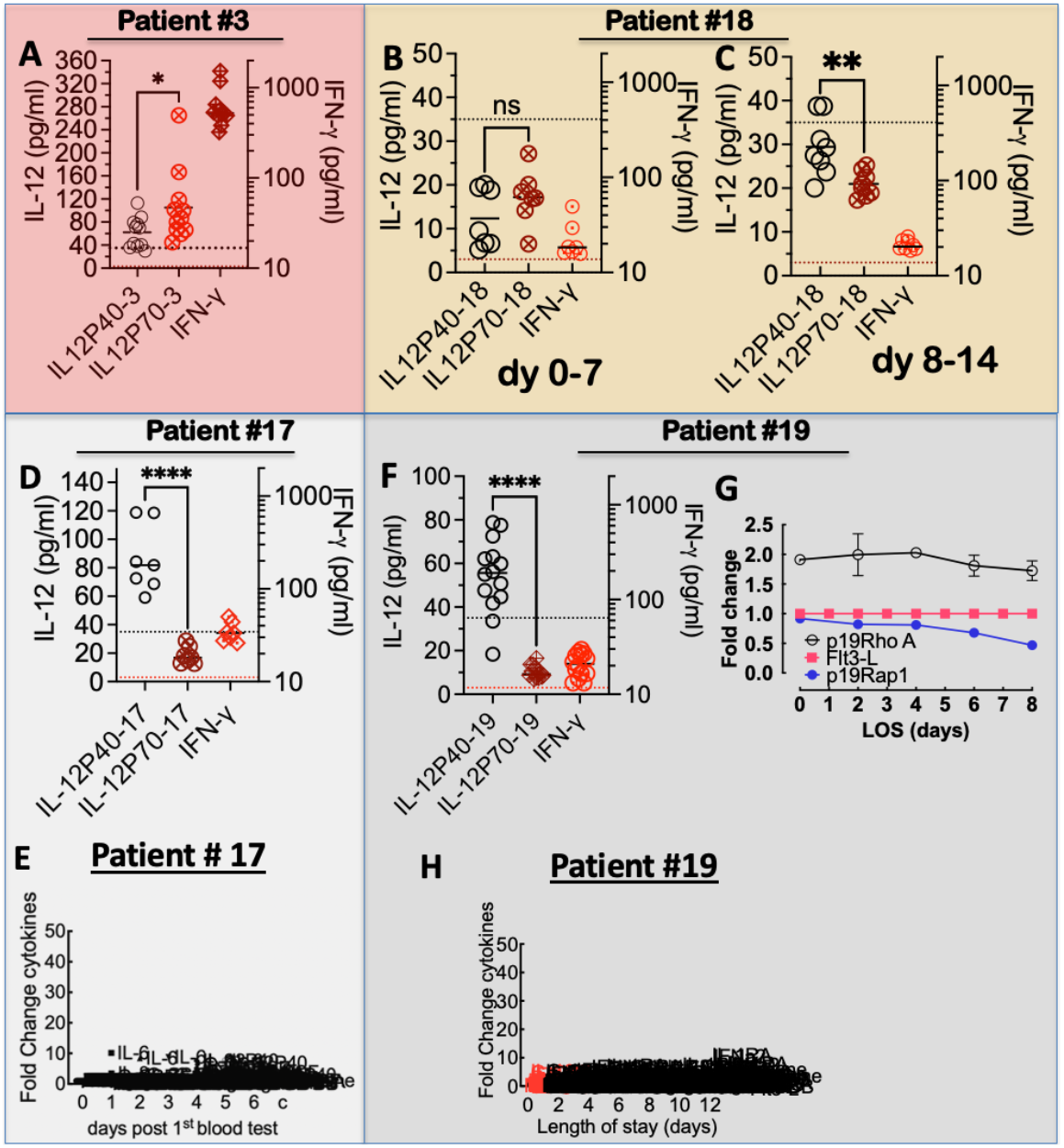
Characteristics of infection relative to sterile inflammation. **A**. Dual plot of IL-12p40, IL-12p70, and IFN-γ for patient #3. Levels of IL-12p40, and IL-12p70 expression were compared using paired t-test; **** *p*< 0.0001, * *p*< 0.05. Evidence of immunosuppression in patient #18’s immune response. **B**. Plot of IL-12p40, IL-12p70, and IFN-γ for patient #18 for days 0-7, showing no significant difference between p40 and p70. **C**. Plot of IL-12p40, IL-12p70, and IFN-γ for patient #18 for days 8-14, showing significant difference between p40 and p70, with no change in IFN-γ expression. Also, overall, IL-12 and IFN-γ expression is comparable to uninfected individuals. Levels of IL-12p40, and IL-12p70 expression were compared using paired t-test; **** *p*< 0.0001, * *p*< 0.05 in GraphPad Prism version 9.3.1. **D**. Plot of IL-12p40, IL-12p70, and IFN-γ for patient 17. Admitted after stab wounds to right neck and chest. ICU LOS was 3 days. No infection was suspected or treated during the enrolled timeframe. Bottom dotted line refers to expression level of IL-12p70 (3.0 pg/ml) and top dotted line refers to IL-12p40 (35pg/ml) for controls. Normal levels of IFN-γ are 15.4 ± 3.8.[75] **E**. Analysis of the longitudinal expression of 41 cytokines (HCYTMAG-60K-PX41 Cytokine kit, Human; Millipore, MA). Data show no immune response. **F**. The plot of IL-12p40, IL-12p70, and IFN-γ for patient 19. The patient was admitted after penetrating trauma to the abdomen. ICU LOS was 7 days. No frank sepsis during the enrolled timeframe. **G**. Analysis of patient #19’s plasma for Rap1·GTP and RhoA·GTP and Flt3-L, showing no immune changes over the test period. H. **E**. Analysis of the longitudinal expression of 41 cytokines (HCYTMAG-60K-PX41 Cytokine kit, Human; Millipore, MA).

